# Validation study of a diffusion MRI derived vessel density biomarker for detecting viral hepatitis-b induced liver fibrosis

**DOI:** 10.1101/633024

**Authors:** Ben-Heng Xiao, Hua Huang, Li-Fei Wang, Shi-Wen Qiu, Sheng-Wen Guo, Yì Xiáng J. Wáng

**Author notes:** Correspondence to: Dr. Yì Xiáng Wáng. Department of Imaging and Interventional Radiology, Faculty of Medicine, The Chinese University of Hong Kong, Shatin, New Territories, Hong Kong SAR., Dr Sheng-Wen Guo. Department of Biomedical Engineering, South China University of Technology, Guangzhou, Guangdong Province, China.

## Abstract

**Aim:** Liver vessel density can be evaluated by an imaging biomarker DDVD (diffusion derived vessel density): DDVD/area(b0b2) = Sb0/ROIarea0 – Sb2/ROIarea2, where Sb0 and Sb2 refer to the liver signal when *b* is 0 or 2 (s/mm^2^); ROIarea0 and ROIarea2 refer to the region-of-interest on *b*= 0 or 2 images; and Sb2 may be replaced by Sb15 (*b*=15). This concept was validated in this study.

**Materials and Methods:** Liver diffusion images were acquired at 1.5T. For a scan-rescan repeatability study of 6 subjects, *b*-values of 0 and 2 were used. The validation study composed of 26 healthy volunteers and 19 consecutive suspected chronic viral hepatitis-b patients, and diffusion images with 16 *b*-values of 0, 2, 4, 7, 10, 15, 20, 30, 46, 60, 72, 100, 150, 200, 400, 600 were acquired. Four patients did not have liver fibrosis, and the rest were four stage-1, three stage-2, four stage 3, and one stage-4 patients respectively.

**Results:** Intraclass correlation coefficient for repeatability was 0.994 for DDVD/area(Sb0Sb2), and 0.978 for DDVD/area(Sb0Sb15). In the validation study, DDVD/area(Sb0Sb2) and area(Sb0Sb15) were 14.80±3.06 and 26.58±3.97 for healthy volunteers, 10.51±1.51 and 20.15±2.21 for stage 1-2 fibrosis patients, and 9.42±0.87 and 19.42±1.89 for stage 3-4 fibrosis patients. For 16 patients where IVIM analysis was performed, a combination of DDVD/area, PF, and Dfast achieved the best differentiation for non-fibrotic livers and fibrotic livers. DDVD/area were weakly correlated with PF or Dfast.

**Conclusion:** Both DDVD/area(Sb0Sb2) and area(Sb0Sb15) are useful imaging biomarker to separate fibrotic and non-fibrotic livers, with fibrotic livers had lower measurements.

---

Chronic liver disease is a major public health problem, accounted for approximately 1.3 million deaths worldwide in 2015 (1). The end result of untreated chronic liver disease is inflammation, loss of liver parenchyma, and healing by fibrosis and regeneration. Earlier stage liver fibrosis is more amenable to therapeutic intervention. In the early stages of fibrosis when the cause has been treated (e.g., hepatitis B or C), regression occurs in at least 70% of patients with the right antiviral management (2, 3). The regression of liver fibrosis can be complete in early stages, whereas partial and prolonged recovery occurs in late or advanced stages (4). Treatment with combined therapies on underline etiology and fibrosis simultaneously might expedite the regression of liver fibrosis and promote liver regeneration. Early detection of liver fibrosis are important for early institution of treatment and assessing potential for regression and prognosis.

Currently there is no established non-invasive diagnostic method to detect and grade early stage liver fibrosis (5). The reference standard for detection and staging of liver fibrosis remains being biopsy; however it is invasive and frequently causes pain and discomfort, with risk of bleeding and hospitalization, subject to sampling errors, and not suitable for longitudinal monitoring. Liver fibrosis is associated with reduced liver perfusion (6-8), and progressive loss of endothelial fenestration and deposition of collagen in the space of Disse. These processes reduce the rate of blood flow and prolong its transit time. On diffusion weighted imaging, blood vessels show high signal when there is no diffusion gradient (*b*=0 s/mm^2^), while show low signal even when very low *b*-values (such as 1 s/mm2) is applied. Recently Wang [9] proposed that liver vessel density can be measured by a diffusion weighted imaging derived surrogate biomarker (DDVD: diffusion derived vessel density):

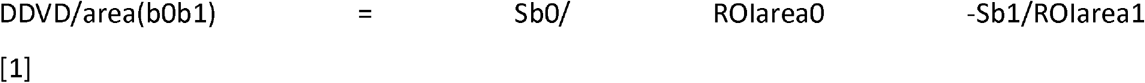

where Sb0 refers to the measured liver signal intensity when *b*=0 s/mm2, and Sb1 refers to the measured liver signal intensity when *b*=1 s/mm^2^. ROIarea0 and Sb1/ROIarea1 refer to the region-of-interest (ROI) on *b*=0 s/mm^2^ and *b*=1 s/mm^2^ images, respectively. Sb1 and ROIarea1 can also be approximated by other low *b*-value diffusion image’s data such as Sb2 which is the measured liver signal intensity when *b*=2 s/mm2. Sb2 may be preferable in cases when Sb1 contain residual high blood signal. Wang [9] further suggested that this surrogate biomarker, DDVD/area, can be used to evaluate the existence and severity of liver fibrosis.

This study aims to develop a method to calculate liver DDVD/area, evaluate their measure-remeasure repeatability, and validate their application in a database composed of healthy livers and fibrotic livers. Moreover, as currently some clinical MRI scanners do not allow low *b*-value less than 10 s/mm^2^ (excerpt *b*=0), this study also evaluated the usefulness of DDVD/area(b0b15)= Sb0/ROIarea0-Sb15/ROIarea15. In addition, the diagnostic performance of a combination of DDVD/area parameter and IVIM (intravoxel incoherent motion) parameters was tested.

## Material and Methods

The MRI data acquisition was approved by the local institutional ethical committee, and the informed consent was obtained for all the subjects. The IVIM type of diffusion scan was based on a single-shot spin-echo type echo-planar sequence using a 1.5-T magnet (Achieva, Philips Healthcare, Best, Netherlands). SPIR technique (Spectral Pre-saturation with Inversion-Recovery) was used for fat suppression. Respiratory-gating was applied in all scan participants and resulted in an average TR of 1600 ms, and the TE was 63 ms. Other parameters included slice thickness =7 mm and inter-slice gap 1mm, matrix= 124×97, FOV =375 mm×302 mm, NEX=2, number of slices =6.

The measure-remeasure repeatability study was performed on six healthy subjects (4 males and 2 females, mean age: 37 yrs, range: 20-58 yrs) during Apr 21 2019 to 19 May 19,2019. For this, volunteers were scanned twice (scan-1 and scan-2) during the same session, with the subjects’ position and selected scan planes unchanged. Images with 16 *b*-values of 0, 2, 4, 7, 10, 15, 20, 30, 46, 60, 72, 100, 150, 200, 400, 600 s/mm^2^ were acquired, and images with b values of 0, 2 s/mm^2^ were used in the current study. The validation study used a database collected during July 27, 2017 to Nov 2, 2018 [10]. There were 26 healthy volunteers (14 males, 12 females, mean age: 24 yrs old; range: 20–41yrs old) and 19 consecutive patients suspected of liver fibrosis with liver biopsy results. Three patients had chronic viral hepatitis-b infection, but biopsy did not show liver fibrosis, and one patient’s biopsy result showed only mild simple steatosis. These four patients were all males, aged 19-57 yrs. The liver fibrosis patients (mean age: 46 yrs, range: 22-62 yrs) had four stage-1 subjects, three stage-2 subjects, four stage 3 subjects, and one stage 4 subject, all with chronic viral hepatitis-b. One patient additionally had hepatocellular carcinoma. Images with *b*-values of 0, 2, 15, 20, 30, 45, 50, 60, 80, 100, 200, 300, 600, 800 s/mm^2^ were acquired.

All data analysis was implemented in MATLAB (MathWorks, Natick, MA, USA). For DDVD/area measurement, only the liver tissue right to the right border of the vertebral body was included (Fig-1). ROI for right liver parenchyma was segmented on the *b*=0 s/mm^2^ image (resulting in ROI area of area0) and the *b*=2 (or 15) s/mm^2^ image (resulting in ROI area of area2 or area15) respectively. Since liver fibrosis mainly affect small/micro-vessels, while in severe fibrosis big vessels may even be dilated [11, 12], this study removed big vessels from analysis. For this ‘bigvessel-pixel removing’ process, on the *b*=0 s/mm^2^ image we evaluated and selected a threshold to remove all visible vessels which are of bright signal; on the *b*=2 s/mm^2^ image, or *b*=15 s/mm^2^ image, we evaluated and selected a threshold to remove all visible vessels which are of ‘signal void’. In some cases it is possible that *b*=2 s/mm^2^ image may contain some residual bright signals as well as bright signal of biliary system, using the same principle as for *b*=0 images, bright signals on *b*=2 s/mm^2^ (or *b*=15) image were also removed. Based on visual assessment, the threshold to remove signal vessels was determined individually for each slices and for each cases. As a general rule, the threshold to remove bright signal vessels was around 35% higher than the mean signals of all pixels in the ROI on *b*=0 images; the threshold to remove signal void vessels was 30% lower than the mean signals of all pixels in the ROI on *b*=2 or 15 images. All results of vessel pixel removal process were visually inspected. The two selected thresholds might have led to slight ‘over-kill’ of some parenchyma pixels, but would have ensured all pixels of pure vessel signal to be removed.

**Fig 1.**
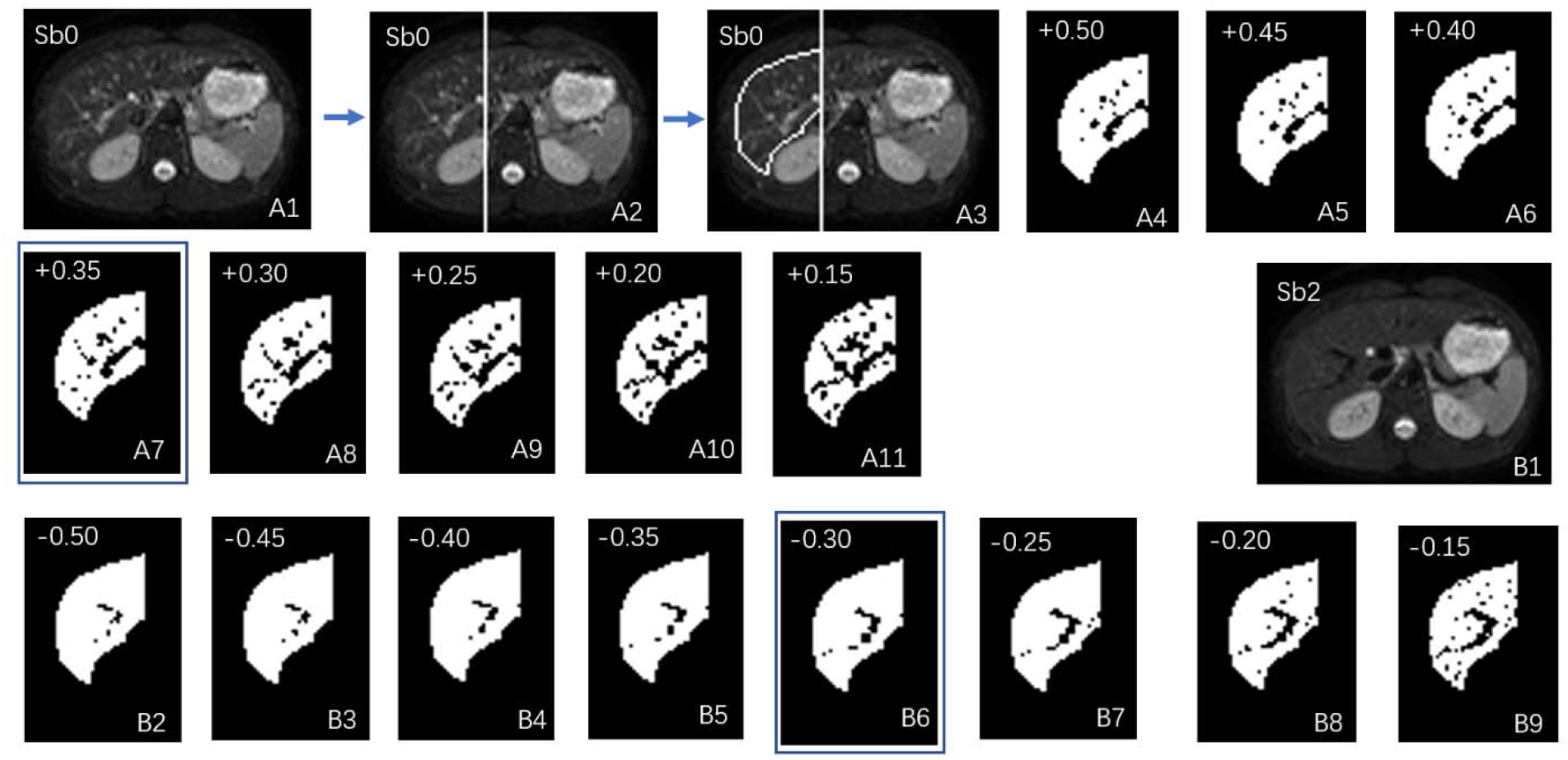
An example of image ‘big-vessel-pixel-removal’ postprocessing of Sb0 image (*b*-value=0 s/mm^2^) and Sb2 image (*b*-value=2 s/mm^2^). A1: the original Sb0 image. A2: a vertical line is drawn along the right border of vertebral body, the liver left to this line is excluded from analysis; A3: the liver right to the vertical line is segmented manually, resulting in an area0. A3: the pixels with signal 50% higher than the mean signal of segmented liver is tentatively excluded; A4: the pixels with signal 45% higher than the mean signal of segmented liver is tentatively excluded. A5-A11 follow the same rule. A7 with the pixels of signal 35% higher than the mean signal excluded shows best results (compromise) in removing ‘bright’ vessel pixels in this case. A8-A11 were considered to have too much ‘over-kill’. B1: the original Sb2 image; B2: the right liver is segmented similar to A3, resulting in an area2, and the pixels with signal 50% lower than the mean signal of segmented liver is excluded. B3-B9 follow the same rule. B6 with the pixels of signal 30% lower than the mean signal of segmented liver excluded show best results (compromise) in removing ‘signal-void ‘ vessel pixels for this case. B7-B9 were considered to have too much ‘over-kill’.

Then two parameters were obtained as

DDVD/area(b0b2) = Sb0/ROIarea0 – Sb2/ROIarea2

DDVD/area(b0b15) = Sb0/ROIarea0 – Sb15/ROIarea15

In this study, the ROI areas of the included slices were broadly similar, the average of DDVDs of individual slice’s was calculated to obtain the value of the liver.

IVIM analysis for PF (perfusion fracture, *f*), Dfast (perfusion related diffusion, *D*\*), and Dslow (true diffusion, *D*) of the study subjects have been presented [10], the IVIM results from our previous study were used for additional analysis in current study. Three patients were excluded for IVIM analysis due to substantial respiratory motion [10]. The *b*=15 s/mm^2^ image was used as the starting point for bi-exponential segmented fitting as detailed previously [9, 10, 13, 14]. The signal value at each *b*-value was normalized by attributing a value of 100 at *b*=15 s/mm^2^ (S_norm_=(SI/SI_15_)×100, where S_norm_ is the normalized signal, SI=signal at a given *b*-value, and SI_15_=signal at *b*=15 s/mm^2^). The threshold was *b*=60 s/mm^2^ were for segmented fitting [14, 15]. For bi-compartmental model, the signal attenuation was modeled according to Eq 1:

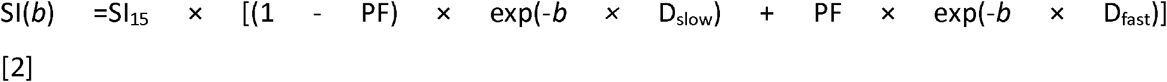

where SI(*b*) and SI_15_ denote the signal intensity acquired with the b-factor value of *b* and *b*=15 s/mm^2^, respectively.

As previously described [13, 15], to estimate the relative distance between the measures of health livers and fibrotic livers, the DDVD/area, PF, Dslow, Dfast values were normalized to range between 0 and 1 according to the formula: (*x_i_* − *x*_min_)/(*x_max_* − *x^min^*).

A 2-D plane was constructed with DDVD/area as the X-axis and PF as the Y-axis. With support vector machine (SVM) approach, the best separating line between health livers and fibrotic livers was defined as *A*X+B*Y+D=0*. In the 2-D coordinate system, the distance between point (*X*_0_, *Y*_0_) and line *A*X+B*Y+D=0* was calculated according to the following equation:

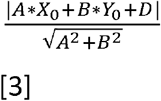

The mean distance of points representing healthy livers to the line and mean distance of points representing fibrotic livers to the line were calculated, and then these two mean distances were added up.

A 3-D coordinate system was then constructed with DDVD/area as the X-axis, PF as the Y-axis, and Dfast or Dslow as the Z-axis. With support vector machine (SVM) approach, the best separating plane between health livers and fibrotic livers was defined as A*X+B*Y+C*Z+D=0. In the 3-d coordinate system, the distance between point (*X*_0_, *Y*_0_, *Z*_0_) and plane A*X+B*Y+C*Z+D=0 was calculated according to following equation:

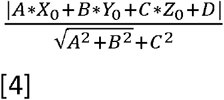

The mean distance of points representing healthy livers to the plane and mean distance of points representing fibrotic livers to the line were calculated, and then these two distances were added up.

For fibrosis patients (n=12), the correlation between DDVD/area vs. Dfast and PF DDVD/area vs. PF were inspected graphically. The values of the three IVIM parameters were re-scaled, with the mean measures of DDVA/area, PF, and Dfast for the fibrosis patients re-scaled to be 1. Moreover, Pearson correlation analysis was performed for correlation between DDVD/area vs. Dfast and PF DDVD/area vs. PF for both all patients (n=16) and volunteers (n=26).

## Results

The scan-rescan repeatability of six healthy volunteers is shown in table 1, intraclass correlation coefficient (ICC) of scan-rescan repeatability was 0.994 and 0.978 for DDVD/area(b0b2) and DDVD/area(b0b15) respectively, suggesting the good scan-rescan repeatability for DDVD/area measurement as well as the robustness of our image post-processing procedure of vessel-pixel removal. However, notable inter-subject variation was also noted with CoV (coefficient of variation) of approximately 0.34 an 0.20 for DDVD/area(b0b2), DDVD/area(b0b15) respectively.

**Tablel-1.** Scan-rescan repeatability measure of six healthy volunteers.

|  | DDVD/area(b0b2) |  | DDVD/area(b0b15) |  |
| --- | --- | --- | --- | --- |
|  | scan 1 | scan 2 | scan 1 | scan 2 |
| subject 1 | 20.2 | 21.3 | 30.8 | 31.7 |
| subject 2 | 11.8 | 11.2 | 22.0 | 22.4 |
| subject 3 | 18.6 | 18.8 | 29.6 | 28.7 |
| subject 4 | 12.3 | 12.8 | 23.6 | 22.2 |
| subject 5 | 9.1 | 9.7 | 19.1 | 20.1 |
| subject 6 | 9.7 | 10.1 | 20.0 | 20.8 |
| mean | 13.6 | 14.0 | 24.2 | 24.3 |
| SD | 4.7 | 4.9 | 4.9 | 4.7 |
| CoV | 0.3428 | 0.3493 | 0.2039 | 0.1947 |

For the validation study, the results of DDVD/area(b0b2) and DDVD/area(b0b15) for healthy volunteers and patients are shown in Fig-2 and Table-2. The mean value of DDVD/area(b0b2) and DDVD/area(b0b15) for the validation study volunteers were are similar to the volunteers of scan-rescan repeatability study (table-1). A trend is seen that fibrotic livers had lower measurement of DDVD/area, and also a severer grade of fibrosis was associated with even lower measurement. On the other hand, DDVD/area of four patients without liver fibrosis showed values similar to healthy livers (Fig-2). Based on the patient/volunteer (pt/vol) ratio, which was the mean measurement for a patient group divided by the mean measurement for healthy volunteers, and also Fig-2, DDVD/area(b0b2) and DDVD/area(b0b15) demonstrated broadly similar performance in separating non-fibrotic vs fibrotic livers, with slightly smaller pt/vol ratio for DDVDb0b2 thus slightly in favor of DDVD/area(b0b2) over DDVD/area(b0b15). Note the smaller the pt/vol ratio, the bigger the difference between the measurements for patients’ value and measurements for healthy volunteers’ value would be.

**Fig-2.**
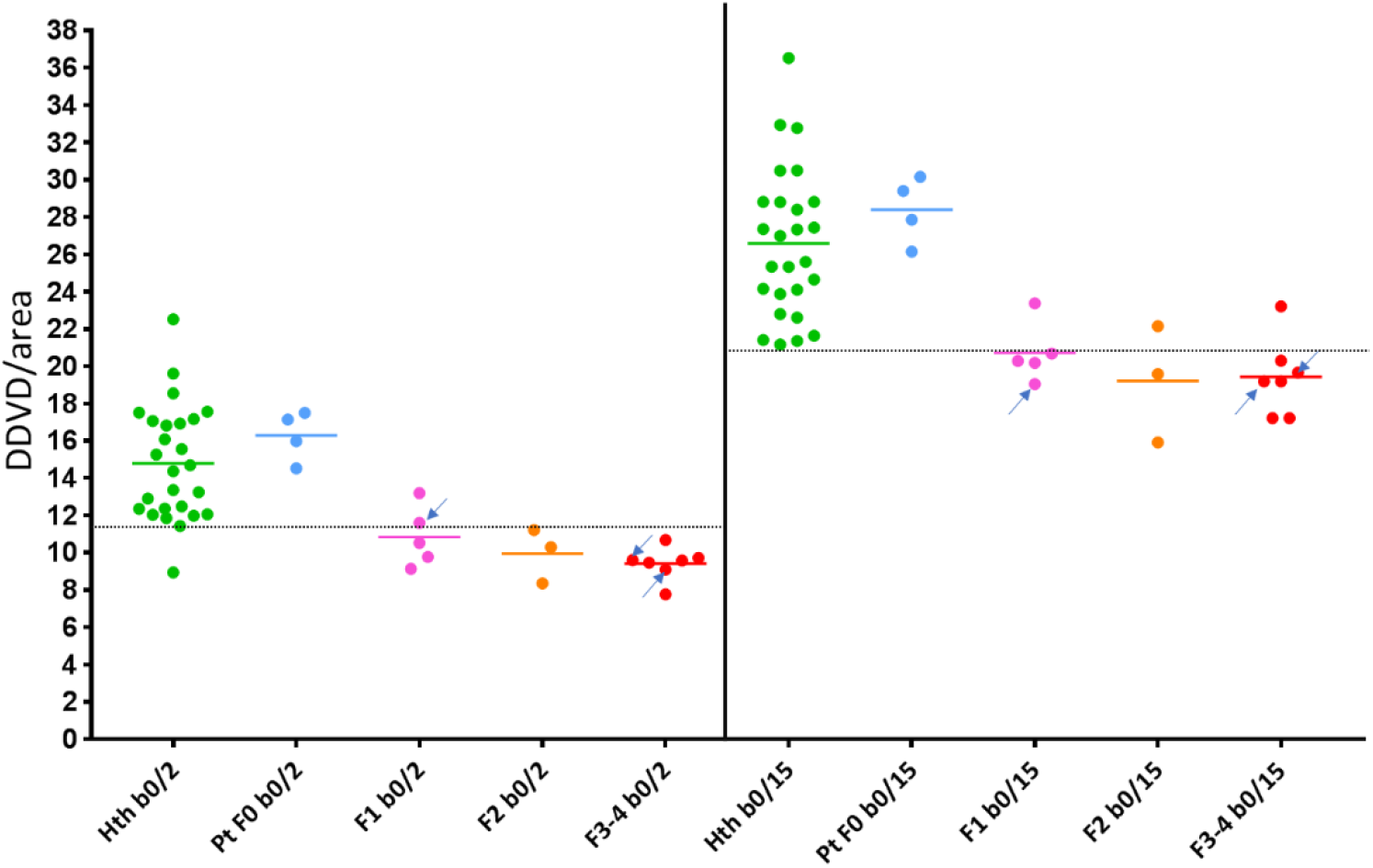
Scatter plot of DDVD/area(b0b2) and DDVD/area(b0b15) measurement of healthy volunteers (Hth, n=26), patients without fibrosis (Pt F0, n=4), and liver fibrosis patients of different stage (F1-F4, n=15). b0/2 denotes measured difference between *b*=0 image and *b*=2 image; b0/15 denotes measured difference between *b*=0 image and *b*=15 image. Arrows denote three patients whose IVIM results could not be measured due to severe respiratory motion (n=3).

**Tablel-2.** Mean and standard deviation of DDVD/area(b0b2) and DDVD/area(b0b15) for healthy volunteers (vol, n=26), patients without fibrosis (pt F0, n=4), and stage-1 and stage-2 liver fibrosis patients (pt F1-2), and stage-3 and stage-4 liver fibrosis patients (pt F3-4). pt/vol ratio (patient/volunteer ratio) = the mean measurement for a patient group divided by the mean measurement for healthy volunteers.

Tablel-1, Scan-rescan repeatability measure of six healthy volunteers.
|  | DDVD/area(b0b2) | DDVD/area(b0b15) |
| --- | --- | --- |
| vol mean ±SD | 14.80±3.06 | 26.58±3.97 |
| vol CoV | 0.2078 | 0.1493 |
| pt F0 mean ±SD | 16.28±1.34 | 28.39±1.78 |
| pt F1-2 mean ±SD | 10.51±1.51 | 20.15±2.21 |
| pt F1-2 pt/vol ratio | 0.7101 | 0.7581 |
| pt F3-4 mean ±SD | 9.42±0.87 | 19.42±1.89 |
| pt F3-4 pt/vol ratio | 0.6365 | 0.7306 |

The 2-D plots of DDVD/area vs PF are shown in Fig 3. If the Y-axis of PF was used as the reference to sperate non-fibrotic livers and fibrotic livers, the healthy volunteers in the orange square which had lower PF measurement (i.e. those close to fibrotic patients in distribution), can be further separated from fibrotic patients by DDVD/area in Y-axis. Furthermore, based on Fig 3, an additional Z-axis was introduced with Dslow or Dfast. The mean relative distances between volunteers’ cluster and patients’ cluster, as calculated with equation-3 and equation-4, are shown in table-3. With the assumption that bigger distance suggests potential better differentiation of non-fibrotic liver and fibrotic livers, table-3 shows the introduction of Dslow or Dfast as a third axis further improved the distance between volunteers’ cluster and patients’ cluster, and even more so with Dfast. Results in table-3 confirm the data in table-2 so that DDVD/area(b0b2) slightly outperformed DDVD/area(b0b15). The 3-D plots of DDVD/area with PF, and Dfast as three axes are shown in Fig-4.

**Fig-3.**
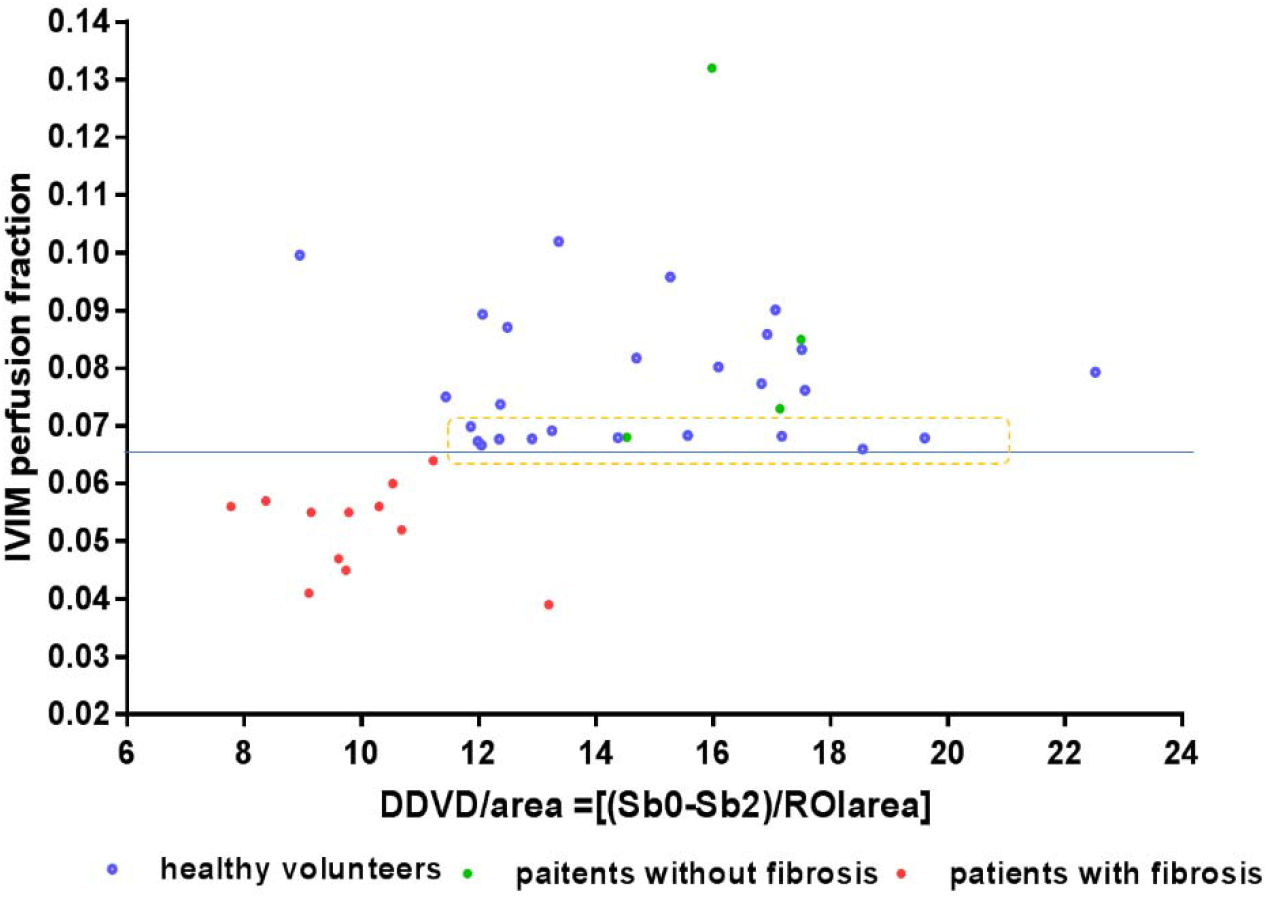
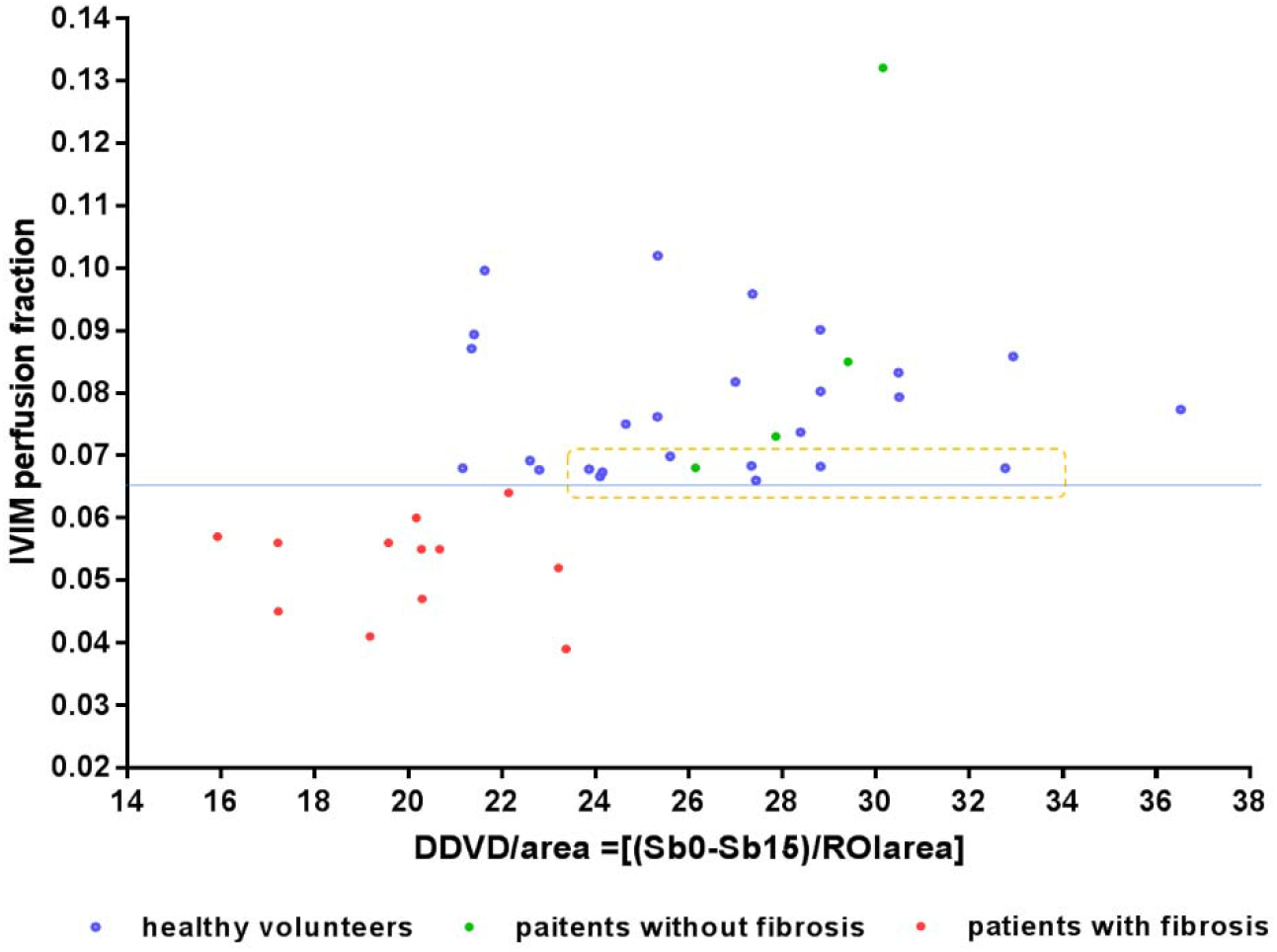
2-D Scatter plot of DDVD/area measurement and PF for of healthy volunteers (blue dot, n=26), patients without fibrosis (green dots, n=4), and liver fibrosis patients (red dots, n=12). Note, compared with Fig-2, there are three patients whose IVIM results could not be measured due to severe respiratory motion, thus these patients are missed in this Figure. (A) for DDVD/area(b0b2) and (B) for DDVD/area(b0b15).

**Figure 4.**
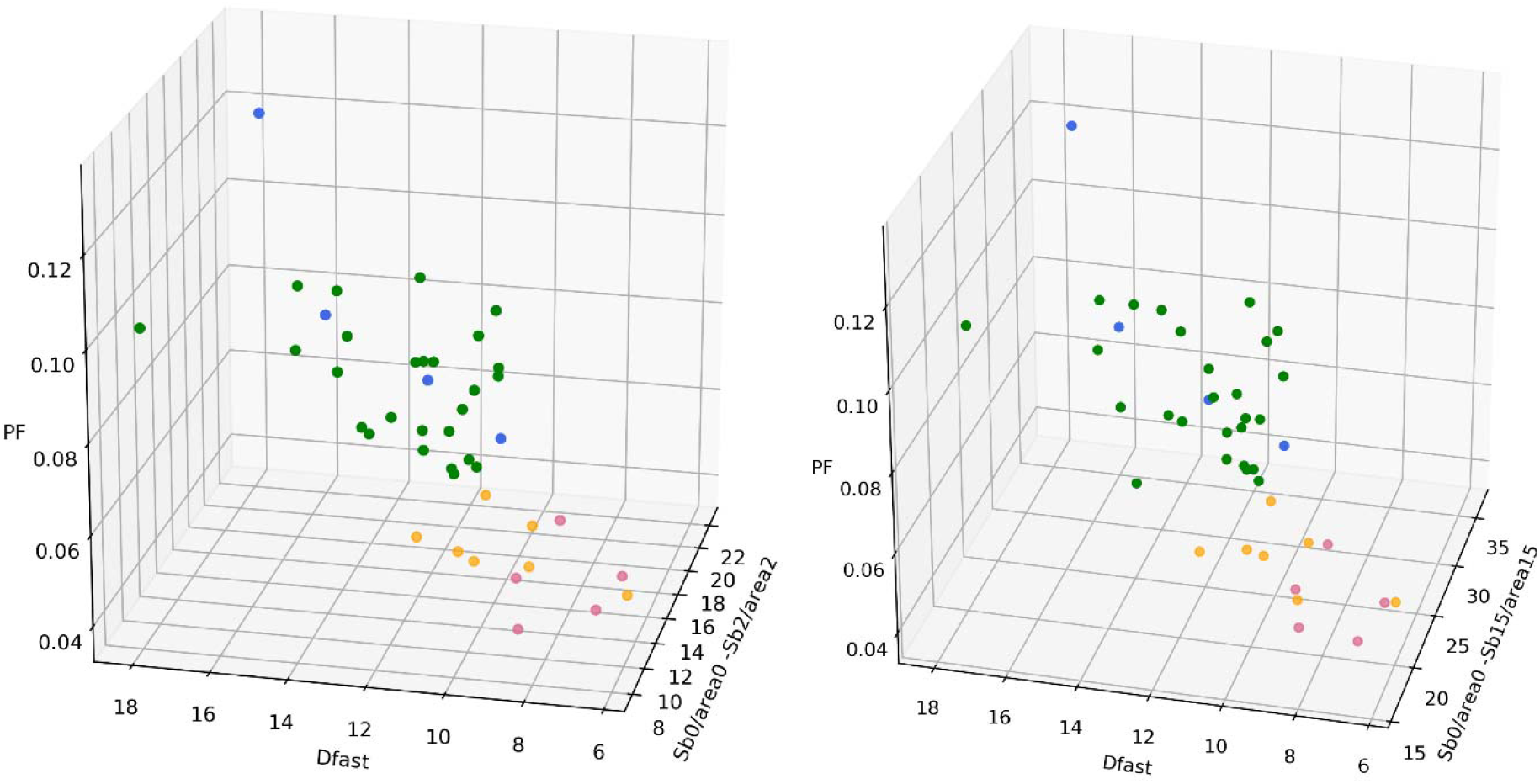
Three-dimensional display of healthy volunteer group (green dots, n=26), patients without liver fibrosis (blue dots, n=4), stage 1–2 liver fibrosis patient group (orange dots, n=7), and stage 3–4 liver fibrosis patient group (red dots, n=5). (A) for DDVD/area(b0b2) and (B) for DDVD/area(b0b15). Each dot represents one participant. The volunteer group and liver fibrosis patient group can be completely separated. The distribution of four patients without liver fibrosis resembles healthy volunteers.

**Table 3.** Mean distance (in relative unit) between healthy volunteer cluster and fibrosis patient cluster. The results were calculated with equation-3 and equation-4.

|  | X-axis= DDVD/area(b0b2)<br>Y-axis=PF | X-axis= DDVD/area(b0b15)<br>Y-axis=PF |
| --- | --- | --- |
| 2d without Z-axis | 0.2162 | 0.2185 |
| 3d Z-axis = Dslow | 0.2515 | 0.2612 |
| 3d Z-axis = Dfast | 0.2957 | 0.2629 |

For liver fibrosis patients, the correlation between DDVD/area vs. Dfast and PF DDVD/area vs. PF are graphically demonstrated in Figure 5. Fig-5 shows DDVD/area larger than 1 could be associated with Dfast or PF smaller than 1, and vice versa, thus no clear correlation pattern could be noted. However, Pearson correlation coefficient *r* for patients and volunteers combined was, respectively, 0.419 (p<0.005), 0.51 (p<0.001), 0.44 (p<0.001), 0.52 (p<0.001), DDVD/area(b0b2) vs. Dfast, DDVD/area(b0b15) vs. Dfast, DDVD/area(b0b2) vs. PF, DDVD/area(b0b15) vs. PF; such the correlations were weak yet significant. These relationships can also be visualized in Fig-3.

**Fig-5.**
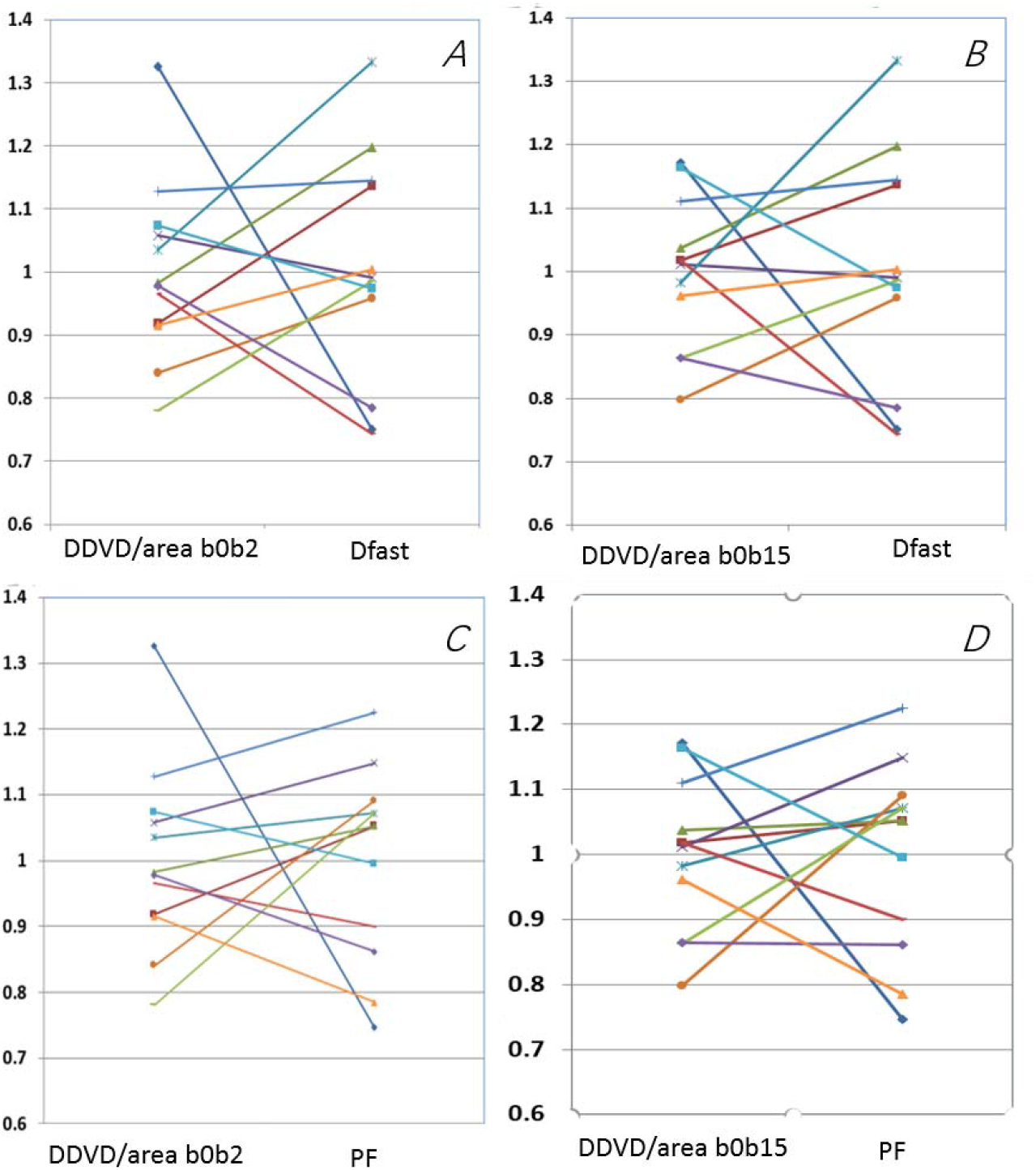
Graphical demonstration of the poor correlation between DDVD/area vs. Dfast and PF DDVD/area vs. PF for fibrotic livers. The mean measures of DDVD/area, PF, and Dfast for the 12 patients with fibrosis were re-scaled to be 1.

## Discussion

Wang et al [9, 13, 15], Huang et al [10], and Li et al [14] recently demonstrated that diffusion MRI derived IVIM measurements can offer high performance in detecting liver fibrosis. However, the disadvantage of IVIM measurement include long scan time to acquire multiple *b*-value imaging data, and this itself is often associated with notable respiration induced motion. Dfast (D*) is also known to be difficult to be fitted precisely [16, 17]. The signal difference between *b*=0 s/mm^2^ image and *b*=1 s/mm^2^ images can be dramatic on diffusion weighted imaging, particularly the vessels show high signal without diffusion gradient while show dark signal when the diffusion gradient is on even at *b*=1 s/mm^2^. This provides the basis for the DDVD/area analysis described in this study. The DDVD/area approaches described in this study offered good scan-rescan repeatability, and provided a useful biomarker for the separation of livers with and without fibrosis, and livers with severe fibrosis tended to have even lower DDVD/area measurement than those with milder liver fibrosis. As could be theorized [9], this study tentatively shows DDVD/area(b0b2) performed slightly better than DDVD/area(b0b15), however, the difference between these two approaches were broadly similar (table-2, table-3). The good scan-rescan repeatability and the similarity between DDVD/area(b0b2)’s results and DDVD/area(b0b15)’s results actually confirmed the robustness of our techniques, particularly the ‘big-vessel-pixel removing’ approach. This may also pave the way for wider application of DDVD concept, as currently many clinical scanners only allow lowest non-zero *b*-value of 10 s/mm^2^. Also of note, the pt/vol ratio for DDVD/area in table-1 is broadly similar to the pt/vol ratio for PF of the IVIM study (table-2 of reference 9), while PF has been proved to be a powerful differentiator of fibrotic livers and non-fibrotic livers [10, 13, 14].

Fibrosis, regenerative nodule formation, and intrahepatic vasoconstriction are classical mechanisms that account for increased intrahepatic vascular resistance in cirrhosis. Mechanisms responsible for the increase in sinusoid resistance include a mechanic factor which is a direct consequence of fibrosis deposition and a dynamic component related to endothelial dysfunction, deficient intrahepatic nitric oxide production, increased vasoconstrictor production, and other factors that promote the increased contraction of hepatic stellate cells [18–20]. As expected, DDVD/area parameter (which is related to micro-vessel density) was shown to be weakly correlated with IVIM perfusion parameter of PF (perfusion fraction) and Dfast (perfusion related fast diffusion). On the other hand, Fig-5 shows for liver fibrosis patients DDVD/area did not have a linear relationship with PF or Dfast, thus DDVD/area PF or Dfast may provide related but in the meantime complementary information. It has been noted that IVIM parameter of Dslow is less sensitive than perfusion parameter of PF and Dfast for detecting liver fibrosis [10, 13, 17]. This study shows the best separation between non-fibrotic livers and fibrotic livers was achieved by a combination of DDVD/area(b0b2), PF, and Dfast (table-3). Using the same method in the current study (equation-4), we additionally quantified the distance when a combination of PF, Dfast, and Dslow was applied, the distance was 0.2488, as opposed to 0.2957 when combination of PF, Dfast, and Dslow DDVD/area(b0b2) was applied (table-3). This difference may be important for separating marginal cases of non-fibrotic and fibrotic liver cases.

The same as IVIM analysis [10], the DDVD/area measures of the three patients of chronic viral hepatitis-b without fibrosis and one patient with simple steatosis resembled those of the healthy volunteers. The diffusion perfusion of yet another 4 patients presented by Li et al [14], further suggest that while pathological process of fibrosis can drive down the liver blood perfusion (as shown with decreased DDVD/area, Dfast and PF), while mere chronic viral hepatitis-b without fibrosis could have normal liver blood perfusion as well as diffusion [10, 14].

In comparison to IVIM analysis, DDVD analysis has the advantage of simplicity, and image data (*b*=0 and *b*=2 or =15) can be acquired by a signal breathhold. In this study, for the three patients excluded for IVIM analysis due to substantial respiratory motion [10], DDVD/area analysis could be performed. In fact, these three patients could be largely separated from heathy volunteers by DDVD/area analysis alone [Fig-2]. As the *b*=2 images were acquired immediately before/after the acquisition of *b*=0 image, thus the respiration induced liver position displacement was usually not substantial. Due to its relative simplicity, DDVD may provide first line testing for assessing liver vessel density. In some cases, when the distribution of patients results is sufficiently different from the known values of healthy subjects, then a diagnostic decision may be made. For cases with ambiguous DDVD results, then additional IVIM scan can be added. It is also likely that a multi-parametric approach will have even better accuracy for evaluating the spectrum of chronic liver disease [21–23], such as IVIM [9, 10, 13, 14, 24, 25], liver T1/T1rho/T2 relaxivity [26–31] and elastography [32, 33] may offer full analysis of liver parenchyma characteristics.

There are a few limitations this study. This a preliminary proof-of-concept study, both data acquisition and data post-processing can be improved in the future. The inter-subject variation between healthy subjects was large in this study. We believe this can be much mitigated by increasing the number of slices scanned, as well as the *b*=0 image and *b*=2 (*b*=15) image scanned by a breathhold technique. As noted, the image data were acquired with respiratory gating in this study, thus the *b*=0 image and *b*=2 (*b*=15) image were not perfectly matched in anatomical location. Currently, the ‘big-vessel-pixel removing’ process was done manually, how to automate this process would be one of our next research priorities. It would be particularly useful if an on-the-spot DDVD/area computing can be performed immediately after the diffusion image data are acquired, so to help select next imaging sequences. All our patients had liver fibrosis due to viral hepatitis-b. Whether results of our study can be generalized to liver fibrosis of other causes, such as NASH (nonalcoholic steatohepatitis), remains to be validated. Our volunteers were on average younger than the patients, so that patients groups and control group were nor matched in age. Finally, we did not control the meal status for this study, as post-meal and fasted status may influence blood flow to the liver, for future studies it can be recommended that patients fast for 6 hours before liver imaging [34].

In conclusion, the DDVD/area approaches described in this study offered good scan-rescan repeatability and was a useful biomarker for the separation of livers with and without fibrosis, and livers with severe fibrosis tended to have even lower DDVD/area measurement than those with milder liver fibrosis. The combined use of DDVD/area and IVIM parameters improved the separation of fibrotic and non-fibrotic livers. Diffusion MRI derived perfusion biomarkers, including DDVD and IVIM, may be able to play important role in liver fibrosis management, both for detection as well as longitudinal monitoring. It can also be expected that the experience learned for liver fibrosis evaluation can also be useful for DDVD analysis of other organs and other pathologies, such as for perfusion-rich tumors.

## Acknowledgement

This work was partially supported by a grant from the Research Grants Council of Hong Kong SAR (Project No. 2141061) and the Sanming Project of Medicine in Shenzhen (SZSM201612053).

